# Parallel neuroinflammatory pathways to cerebrovascular injury and amyloid-beta in Alzheimer’s disease

**DOI:** 10.1101/2024.10.03.616579

**Authors:** Batool Rizvi, Jenna N. Adams, Alison Bamford, Soyun Kim, Mithra Sathishkumar, Nicholas J. Tustison, Lisa Taylor, Nandita Tuteja, Liv McMillan, Bin Nan, Hengrui Cai, Yuritza Y. Escalante, Novelle Meza, Alyssa L. Harris, Rond Malhas, Adam M. Brickman, Mark Mapstone, Elizabeth A. Thomas, Michael A. Yassa

**Affiliations:** Center for the Neurobiology of Learning and Memory, University of California, Irvine, CA, USA; Department of Neurobiology and Behavior, University of California, Irvine, CA, USA; Institute for Interdisciplinary Salivary Bioscience Research, University of California, Irvine, CA, USA; Department of Radiology and Medical Imaging, University of Virginia, Charlottesville, VA, USA; Department of Epidemiology and Biostatistics, University of California, Irvine, CA, USA; Department of Statistics, University of California, Irvine, CA, USA; Department of Neurology, University of California, Irvine, CA, USA; Taub Institute for Research on Alzheimer’s Disease and the Aging Brain, College of Physicians and Surgeons, Columbia University, New York, NY, USA; Gertrude H. Sergievsky Center, College of Physicians and Surgeons, Columbia University, New York, NY, USA; Department of Neurology, College of Physicians and Surgeons, Columbia University, New York, NY, USA; Department of Neurosciences, The Scripps Research Institute, La Jolla, CA, USA

**Author notes:** **Corresponding author**s: Michael A. Yassa, Ph.D., 1418 Biological Sciences 3, Irvine CA, 92697-4550 | |, Batool Rizvi, M.S., 1400 Biological Sciences 3, Irvine CA, 92697-4550 | |.

## Abstract

**Importance:** While the hallmark pathologies of amyloid-beta (Aβ) and tau in Alzheimer’s disease (AD) are well documented and even part of the definition, upstream neuroinflammation is thought to play an important role but remains poorly understood.

**Objectives:** We tested whether two distinct neuroinflammatory markers are associated with cerebrovascular injury and Aβ, and whether these markers are associated with plasma phosphorylated tau (pTau) concentration, medial temporal lobe (MTL) cortical and hippocampal atrophy, and memory deficits. We examined neuroinflammatory markers plasma YKL-40 and GFAP, due to previous conflicting evidence relating YKL-40 and GFAP to AD pathogenic markers.

**Design:** Cross-sectional data from a community observational study (Biomarker Exploration in Aging, Cognition, and Neurodegeneration - BEACoN) were included.

**Setting:** All participants were enrolled in a single site, at University of California, Irvine.

**Participants:** 126 participants were included if they had at least one of the following measures available: neuropsychological data, MRI, Aβ-PET, or plasma.

**Exposures:** Plasma YKL-40 and plasma glial fibrillary acidic protein (GFAP) levels.

**Main outcomes and measures:** White matter hyperintensity (WMH) volume, 18F-florbetapir (FBP) PET mean SUVR, plasma phosphorylated tau (pTau-217) concentration, MTL cortical thickness, hippocampal volume, and memory function assessed by Rey Auditory Verbal Learning Test. Using path analysis, we tested whether higher plasma YKL-40 and GFAP are associated with WMH and Aβ, and whether these converge to downstream markers of tauopathy, MTL neurodegeneration, and memory deficits.

**Results:** In older adults without dementia (N=126, age=70.60±6.29, 62% women), we found that higher plasma YKL-40 concentration was associated with greater WMH volume, while higher plasma GFAP concentration was related to increased FBP SUVR. Further, higher plasma GFAP, WMH and FBP SUVR were independently associated with increased pTau-217. In turn, plasma pTau-217 was associated with reduced MTL cortical thickness and hippocampal volume. Subsequently, only reduced hippocampal volume was related to lower memory function.

**Conclusions and Relevance:** Neuroinflammatory markers contribute to parallel pathways of cerebrovascular injury and Aβ, which converge to tau-associated neurodegeneration and memory deficits in older adults. These observations underscore the need for a more comprehensive approach to developing an AD framework and treatment strategies.

**KEY POINTS:** *Question:* How does neuroinflammation impact downstream features of cerebrovascular injury and amyloid-beta (Aβ) in Alzheimer’s disease?

*Findings:* In this study of 126 older adults without dementia, we found evidence for two distinct neuroinflammatory pathways that lead to neurodegeneration and memory deficits. One path involves plasma YKL-40 and its impact on cerebrovascular injury, as measured by white matter hyperintensities (WMH) on MRI scans. The other involves plasma glial fibrillary acidic protein (GFAP) and its impact on Aβ deposition measured via 18F-florbetapir (FBP) PET. Both pathways converged on tauopathy, measured by plasma pTau-217, which was associated with lower medial temporal lobe (MTL) cortical thickness and hippocampal volume, and consequently, memory deficits.

*Meaning:* Inflammation acts on Alzheimer’s disease mechanisms via multiple distinct and parallel pathways which converge downstream onto neurodegeneration.

**Graphical Abstract:** 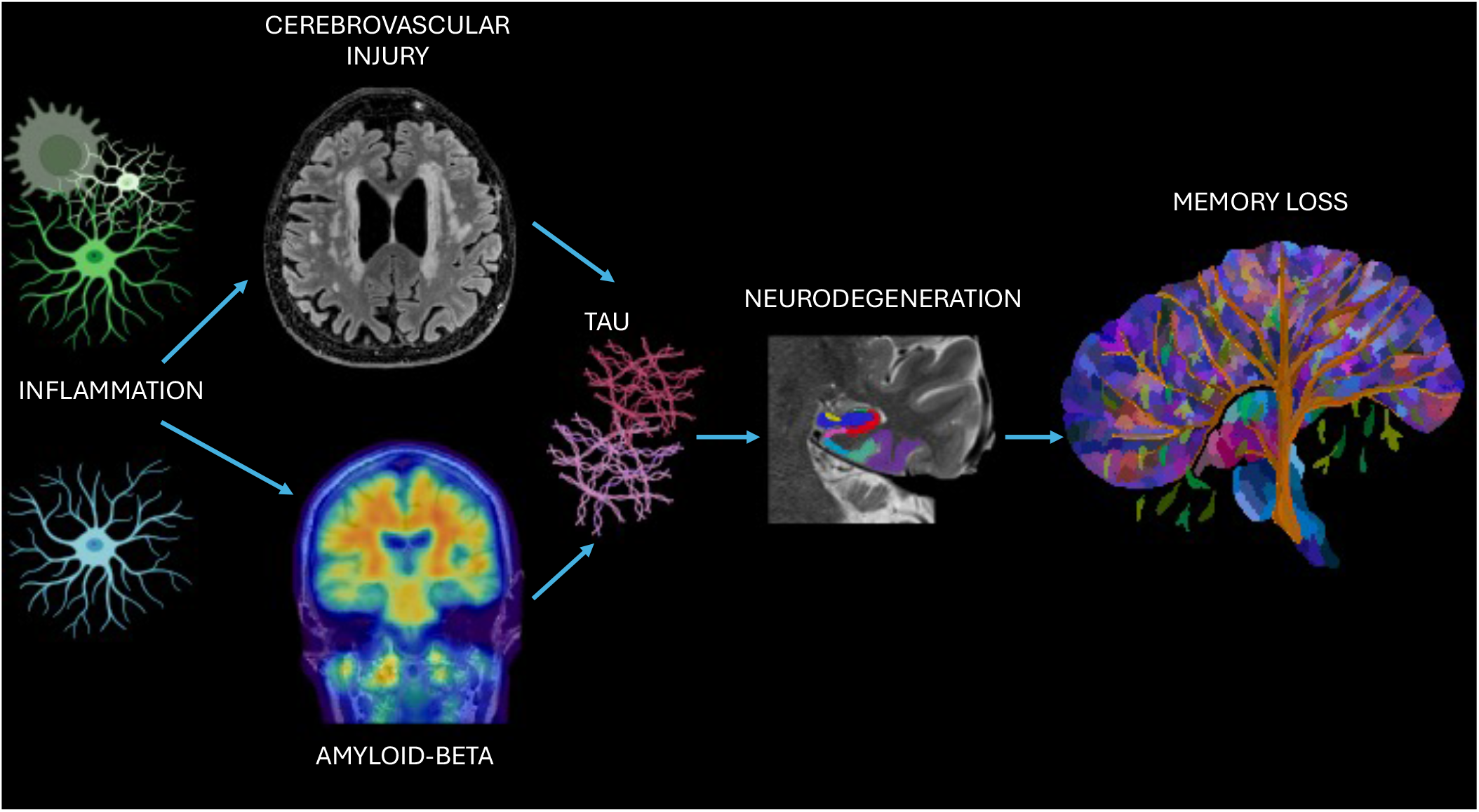

**Credit:** BioRender was used to help create this graphical abstract.

## 1. INTRODUCTION

Alzheimer’s disease (AD) is defined by two pathologies – amyloid-beta (Aβ) plaques and neurofibrillary tangles ^1^. However, less emphasis is placed on the heterogeneity of processes in AD, which impedes us from determining novel potential mechanistic pathways preceding clinical symptoms and identifying effective markers for treatment. While Aβ is hypothesized to initiate a biological cascade that leads to accelerated deposition and spread of tau pathology and neurodegeneration ^2-4^, there are likely multiple pathways that converge to promote neurodegeneration and cognitive impairment. One such pathway involves the interplay between neuroinflammation and cerebrovascular injury, which may parallel the cascading impact of Aβ pathology ^3,5^.

Reactive astrocytes and microglia are important to both Aβ and tau pathogenesis ^6,7^ and the possible bidirectional roles of neuroinflammation with AD proteinopathy have been examined ^8,9^. For example, initial glial activation is thought to be an early mechanism to eliminate Aβ, yet this process often leads to a vicious cycle of development of further chronic neuroinflammatory and neurotoxic damage ^10^. Specific reactive astrocytic and microglial markers, such as glial fibrillary acidic protein (GFAP) and chitinase-3-like protein 1 (YKL-40), contribute to higher risk of AD ^11-14^. YKL-40 has emerged as an important candidate biomarker for examining the clinical progression of AD ^15^, and can be detected at the earliest stage of pathogenesis ^16-18^. Similarly, GFAP is a valuable marker in the early detection and prediction of the course of AD ^19^. One study comparing several candidate AD plasma biomarkers found that plasma GFAP combined with AD risk factors had the highest accuracy in differentiating between cognitively unimpaired older adults with amyloidosis and without amyloidosis, demonstrating its potential as a diagnostic marker in AD ^11^. Comparing whether GFAP and YKL-40 are differentially linked to Aβ and cerebrovascular injury has not yet been investigated. One marker reflecting cerebrovascular injury and small vessel cerebrovascular disease, broadly studied in aging and dementia ^20-22^, are white matter hyperintensities (WMH), which are visualized as areas with increased brightness on T2-weighted magnetic resonance imaging (MRI).

One study found that CSF GFAP was related to Aβ while CSF YKL-40 was related to higher tau-PET ^23^. Ferrari-Souza et al. ^23^ further found that CSF GFAP and YKL-40 mediated the effects of Aβ and tau respectively on hippocampal atrophy, which was subsequently related to worse cognition. Similarly, Pelkmans et al. ^24^ reported CSF Aβ is linked to GFAP. However, Pelkmans et al. ^24^ determined that CSF YKL-40 mediated the effect of Aβ on tau-phosphorylation and its association to neurodegeneration. Furthermore, another study suggested that GFAP mediated the association between Aβ and tau ^25^. Increases in plasma GFAP levels have also been associated with WMH in early-onset AD, late-onset AD, and in adults with Down syndrome (DS) ^26,27^. Similarly, there is growing evidence of the association between YKL-40 and WMH ^28^. A longitudinal study demonstrated a strong association between plasma and CSF YKL-40 levels ^14^. Another study indicated that plasma GFAP was a sensitive biomarker that outperformed CSF GFAP in detecting amyloid pathology in early AD ^29^. Given the inconsistency of the reported links of GFAP and YKL-40 with Aβ, tau, and WMH, we tested these associations in the context of an AD pathogenic pathway.

A novel feature of the current study is the examination of cerebrovascular injury and its interface with neuroinflammation, Aβ, and tau burden. Past work indicates an association of WMH with structural imaging markers of medial temporal lobe (MTL) neurodegeneration and memory deficits in older adults ^30,31^. Furthermore, related work shows that WMH precede and promote tau ^32,33^. We hypothesize that one mechanism by which WMH lead to MTL and hippocampal neurodegeneration is through tau burden. Overall, we propose that cerebrovascular injury and dysfunction is a distinguishable pathology from amyloidosis. We further suggest that these two types of pathologies may be associated with different neuroinflammatory cascades. This is in contrast with the single pathogenic pathway proposed by the A/T/N framework ^34,35^.

## 2. MATERIALS AND METHODS

### 2.1 Participants

One-hundred twenty-six community-dwelling adults were included in the study, as part of the Biomarker Exploration in Aging, Cognition, and Neurodegeneration (BEACoN) study. Inclusion criteria for BEACoN consists of age ≥ 60 years and performance on cognitive assessments within age-adjusted normal range (within 1.5 standard deviations). Exclusion criteria includes diagnosis of dementia or mild cognitive impairment, major health problems (e.g. uncontrolled diabetes mellitus, uncontrolled hypertension, history of stroke), comorbid neurological disease, significant psychiatric disorders, use of medication for anxiety or depression or illicit drugs, and MRI or PET contraindications. All participants gave written informed consent and were compensated for their participation. Study procedures were approved by the Institutional Review Board of the University of California, Irvine. The study is in accordance with *The Code of Ethics of the World Medical Association (Declaration of Helsinki)*. Participants were included in the study if MRI, Aβ PET, plasma, or neuropsychological data of interest in our primary analyses were available. Demographic characteristics of the participants, including the sample size of each marker, are included in Table 1.

**Table 1.** Demographic Characteristics and Summary Variables.

| Characteristic | Values |  |
| --- | --- | --- |
| *N* | 126 |  |
| Age, mean (SD) | 70.60 (6.29) |  |
| Education, mean (SD) | 16.57 (2.32) |  |
| Sex/Gender, n (%) women | 78 (61.90%) |  |
| Race and Ethnicity, n (%) |  |  |
| White | 102 (80.95%) |  |
| Asian | 18 (14.29%) |  |
| Black | 1 (0.79%) |  |
| More than 1 race or Other | 5 (3.97%) |  |
| Hispanic | 6 (4.76%) |  |
| Markers (Plasma, Imaging, Neuropsychological) | Raw values | Log-transformed values |
| YKL-40 pg/mL (n=69), mean (SD) | 51.41 (41.23) | 3.73 (0.64) |
| GFAP pg/mL (n=69), mean (SD) | 353.90 (331.89) | 5.65 (0.63) |
| Total WMH, cm<sup>3</sup> (n=112), mean (SD) | 3.60 (8.14) | 1.02 (0.83) |
| Global FBP SUVR (n=108), mean (SD) | 1.11 (0.17) | 0.74 (0.08) |
| pTau-217 pg/mL (n=64), mean (SD) | 0.12 (0.10) | 0.11 (0.08) |
| MTL Cortical Thickness, mm (n=122), mean (SD) | 1.20 (0.06) | 0.79 (0.27) |
| Hippocampal Volume, TIV adjusted (n=122), mean (SD) | 1.96 (0.26) | 1.08 (0.09) |
| Retroactive Interference (n=111), mean (SD) | 0.84 (0.18) | 0.60 (0.10) |
| Delayed Recall (n=111), mean (SD) | 10.72 (3.52) | 2.39 (0.47) |

### 2.2 Magnetic Resonance Imaging

Magnetic resonance imaging (MRI) data were acquired on a 3.0 Tesla Siemens Prisma scanner at the Facility for Imaging and Brain Research (FIBRE) at the University of California, Irvine. The following scans were acquired: Structural T1-weighted MPRAGE (resolution = 0.8 × 0.8 × 0.8 mm, repetition time = 2300 ms, echo time = 2.38 ms, FOV read = 256 mm, slices = 240, slice orientation = sagittal), T2-weighted fluid-attenuated inversion recovery (FLAIR; resolution = 1.0 × 1.0 × 1.2 mm, repetition time = 4800 ms, echo time = 441 ms, FOV read = 256 mm, slices = 160, inversion time = 1550 ms, slice orientation = sagittal) and T2-Turbo Spin Echo (resolution = 0.4 mm × 0.4 mm × 2.0, repetition time = 5000 ms, echo time = 84 ms, FOV read = 190 mm, acquired perpendicular to the long axis of the hippocampus).

### 2.3 MRI analyses

#### 2.3.1 White Matter Hyperintensities Segmentation

Image processing leveraged the open-source ANTsX software ecosystem with a particular focus on specific deep-learning applications developed for neuroimaging made available for both Python and R via the ANTsXNet (ANTsPyNet/ANTsRNet) libraries. Specifically, for the work described here, WMH segmentation used the ANTsPyNet function: sysu_white_matter_hypterintensity_segmentation.

The methods used in this paper for WMH segmentation (see Figure 1A, 1B) and quantification are previously described ^31^. Briefly, a preprocessing scheme is used that includes thresholding for brain extraction and an ensemble (n=3) of randomly initialized 2-D U-nets to produce the probabilistic output, implemented in ANTsXNET ^36^. All WMH masks resulting from the above automated segmentation procedure were manually edited by an expert rater (B.R.) for improved accuracy. These edited global WMH volumes were used in our analyses.

**Figure 1.**
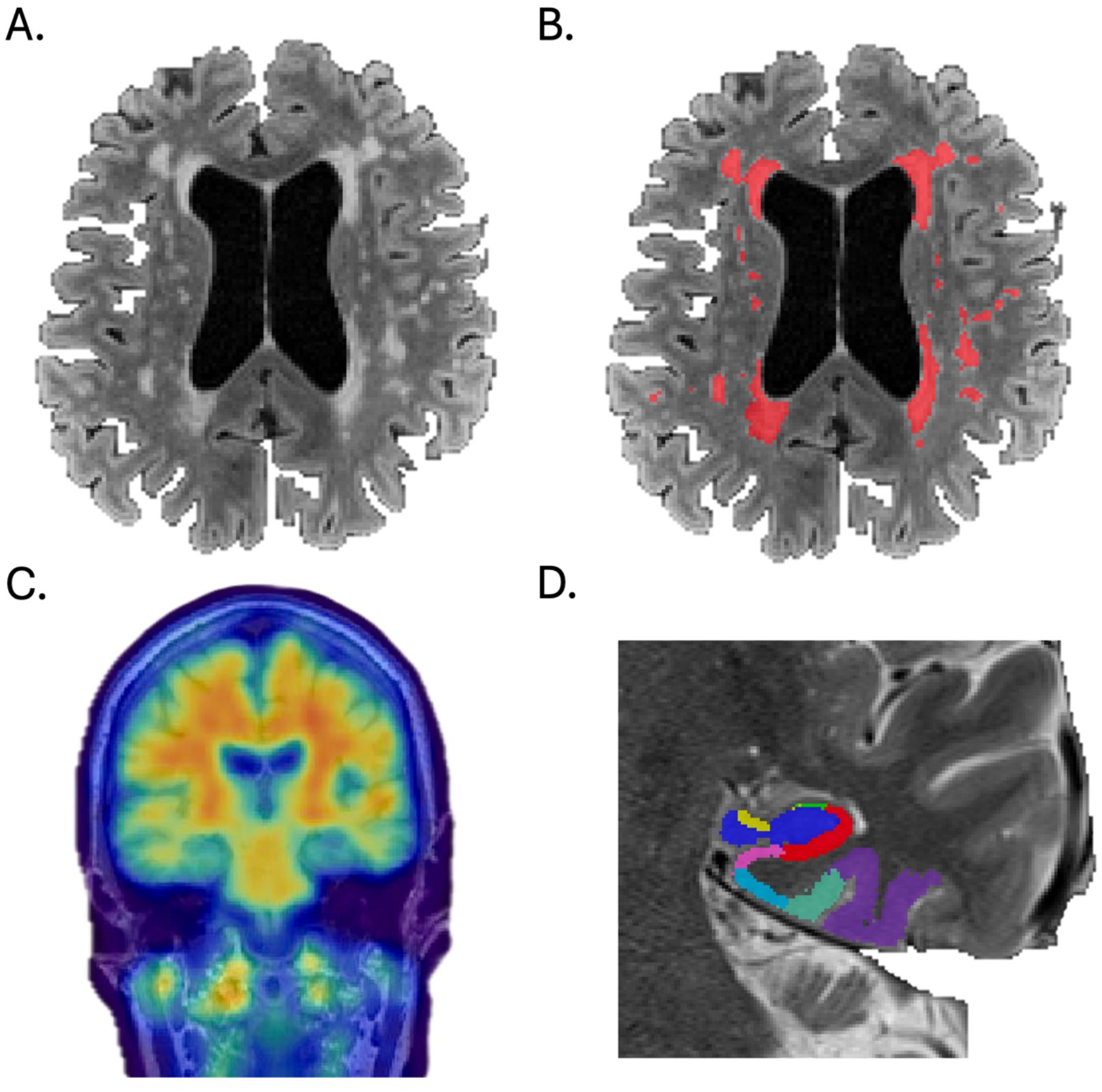
Visualizing imaging analyses A) Unlabeled WMH on an axial T2-FLAIR image. B) Labeled WMH in red on an axial T2-FLAIR image. C) Global FBP SUVR image D) T2 image with hippocampal and MTL cortical subregions labeled using ASHS.

#### 2.3.2 Medial Temporal Lobe Subregion Segmentation

Medial temporal lobe (MTL) subregions were automatically segmented (see Figure 1D) with the Automatic Segmentation of Hippocampal Subfields (ASHS) ^37^ software using T1- and T2-weighted images. The ASHS pipeline implements joint label fusion and corrective learning to segment hippocampal subfields and medial temporal lobe cortical subregions. The left and right MTL segmentation images from ASHS were then loaded using Advanced Normalization Tools software (ANTs). Label geometry measures, including volume and surface area, were extracted from the segmented subregions. For each labeled subregion, the thickness was computed by dividing the volume by the surface area.

The resulting output included thickness of the following MTL cortical subregions in native T2 space: entorhinal cortex (ERC), perirhinal cortex subdivided into Brodmann Areas 35 and 36 (PRC; BA35, BA36), and parahippocampal cortex (PHC). Total MTL cortical thickness was derived from averaging the thickness of these four subregions. Separately, the output included volumes of the following hippocampal subfields: dentate gyrus (DG), CA1, CA2, CA3, and subiculum. All hippocampal subfield volumes were adjusted for intracranial volume (ICV) by dividing the subfield volume by the individual’s ICV. Total hippocampal volume was derived by taking the sum of the ICV-adjusted bilateral hippocampal subfield volumes. Total hippocampal volume was then multiplied by 1000.

### 2.4 Plasma Markers

#### 2.4.1 Plasma Collection

Blood was collected without regard to prandial state, time of day or medication timing. Blood was drawn via venipuncture from each participant into 7 mL lavender top EDTA tubes (BD 366450). Immediately after collection, each tube was gently mixed by inverting 8 to 10 times to ensure proper mixing of blood and anticoagulant, and then placed on wet ice. Blood samples were centrifuged in a swinging rotor bucket within 1 hour of collection at 2600 x RPM at 20°C for 10 minutes. The isolated plasma was transferred and pooled into a sterile 50 mL polypropylene conical tube and mixed by inversion a few times. The plasma samples were aliquoted by 0.750 mL increments into 2 mL polypropylene cryovials. The plasma aliquots were transferred into a −80 °C freezer for storage until required for analysis.

#### 2.4.2 Plasma Biomarker Quantification

Levels of YKL-40 were quantified in plasma samples from BEACoN participants using the U-PLEX Human YKL-40 ECL immunoassay MesoScale Discovery (MSD; Cat #K151VLK Gaithersburg, MD). Levels of GFAP were quantified using the R-PLEX Neurology Panel 1 ^38 K15639^. Assays were run according to MSD manufacturers protocol using plasma samples diluted 1:2 in Diluent 3 or 64 (MSD) for YKL-40 and GFAP, respectively. Samples were assayed after a single thaw to room temperature. Measurements were performed in duplicate, and sample measurements accepted if signal coefficients of variation (CVs) across duplicates were less than 15%. The human YKL-40 and GFAP calibrators provided with the kit were used for generating the standard curve, and sample concentrations (pg/ml) were determined with MSD Discovery Workbench Software using curve fit models. Lower limits of detection (LLoD) were as follows: YKL-40 (0.858 pg/mL) and GFAP (29.1 pg/mL). Plasma p-Tau 217 levels were quantified using an immunoassay developed by ALZpath, using a previously described method ^39^.

### 2.5 Amyloid PET

#### 2.5.1 PET Acquisition

18F-Florbetapir (FBP) positron emission tomography (PET) was used to measure Aβ plaque pathology. PET was acquired on an ECAT High Resolution Research Tomograph (HRRT, CTI/Siemens, Knoxville, TN, USA). Ten mCi of tracer was injected, and then four 5-minute frames were collected from 50-70 minutes post-injection. FBP-PET data was reconstructed with attenuation correction, scatter correction, and 2mm3 Gaussian smoothing.

#### 2.5.2 PET Processing

FBP frame data were realigned, averaged, and then coregistered to the T1-weighted MPRAGE image. T1 MPRAGE images were processed with FreeSurfer v.6.0 to generate native-space regions of interest for PET reference regions and quantification. FBP images were then normalized by a whole cerebellum reference region to produce standardized uptake value ratio (SUVR) images (see Figure 1C). Additional 6mm3 smoothing was then applied to achieve an effective resolution of 8mm3. The mean SUVR of a validated cortical composite region ^40,41^ was then quantified to obtain a global measure of FBP uptake for analysis.

### 2.6 Neuropsychological Testing

Participants were administered the Rey Auditory Verbal Learning Test (RAVLT), which assesses word list learning, immediate and delayed memory ^42^. Our primary outcome measure of memory in this study was retroactive interference (RI) from RAVLT, which reflects resistance to interference during consolidation. RI compares memory for the target list of words before (A5) and after (A6) a distractor list is presented (A6/A5; with lower scores indicating increased interference). Previous work suggests that RI is a more sensitive behavioral biomarker of subtle memory deficits than delayed recall, reflecting AD pathology and MTL subregional integrity ^43,44^. In a supplementary analysis, we used delayed recall from RAVLT (A7), which tests free recall for the target word list after a 20 minute delay from the last learning trial.

### 2.7. Statistical Analysis

Statistical analyses were conducted in R (version 4.3.0) through RStudio. All imaging, plasma, and neuropsychological markers of interest were log-transformed in order to reduce the skewness of the data prior to running path analysis, with a constant of 1 added before transforming to handle any zeros and to preserve non-negative values.

We conducted path analysis, which is a subset of structural equation modeling, using the *lavaan* package in R in a series of linear regressions simultaneously. We used full information maximum likelihood to handle missing data. Independent variables were considered as random. In our path analysis, we tested whether neuroinflammatory markers, including plasma GFAP and YKL-40, are associated with global WMH volume and global FBP SUVR. WMH volume and global FBP SUVR (Aβ) were then tested as two dissociable markers associated with plasma pTau-217. We also tested whether plasma pTau-217 is associated with MTL cortical thickness and hippocampal volume, and subsequently to lower performance on retroactive interference from RAVLT. In the supplementary results, we additionally report results using RAVLT delayed recall as an alternative outcome measure. We assessed goodness of model fit as “acceptable” if: *X*^*2*^ *p* > 0.05, Comparative Fit Index (CFI ≥ 0.90), Tucker-Lewis Index (TLI ≥ 0.90), Root Mean Square Error of Approximation (RMSEA < 0.08), and Standardized Root Mean Square Residual (SRMR < 0.08). Age was not controlled for in the path analysis, as the biomarkers and their relationships being studied are largely age-dependent. Furthermore, as this sample does not not include participants with frank clinical impairment or dementia, adjusting for age would limit our ability to capture important associations among biomarkers and between biomarkers and cognitive function.

## 3. RESULTS

### Path Analysis

In our path analysis model, we tested associations among markers and hypothesized a theoretically driven mechanistic cascade. The overall model fit was considered acceptable: *X*^*2*^ (9, *N*=126) = 21.049, *p* = 0.10, CFI = 0.90, TLI = 0.81, RMSEA = 0.063, and SRMR = 0.068. Plasma YKL-40 levels were not correlated with plasma GFAP levels (see Table 2, Figure 2). Higher plasma YKL-40 levels were associated with greater global WMH, but not with global FBP SUVR. Meanwhile, higher plasma GFAP levels were associated with higher global FBP SUVR, but not with global WMH. Higher plasma GFAP levels were also related to higher plasma pTau-217 levels. Both greater global WMH and increased global FBP SUVR were independently associated with higher plasma pTau-217 levels. Higher plasma pTau-217 levels were in turn associated with both reduced MTL cortical thickness and lower hippocampal volume. Subsequently, only hippocampal volume was related to lower performance on retroactive interference (RAVLT). In a supplemental analysis, we found similar patterns of associations among biomarkers, and determined that lower hippocampal volume was related to lower scores on delayed recall (RAVLT; see Supplemental Table 1). Simple associations among markers included in the path model can be visualized in the scatterplots shown (see Figure 3).

**Table 2.** Results of path analysis.

| Dependent variable | Independent variable | *B* | Standardized Estimate | p-value | 95% CI |
| --- | --- | --- | --- | --- | --- |
| Total WMH (log) ~ | YKL-40 (log) | 0.393 | 0.299 | 0.005 | (0.12, 0.666) |
|  | GFAP (log) | 0.098 | 0.076 | 0.465 | (-0.166, 0.362) |
| Global FBP SUVR (log) ~ | YKL-40 (log) | -0.02 | -0.161 | 0.176 | (-0.05, 0.009) |
|  | GFAP (log) | 0.054 | 0.435 | <0.001 | (0.027, 0.08) |
| pTau-217 (log) ~ | Total WMH (log) | 0.023 | 0.226 | 0.014 | (0.004, 0.041) |
|  | Global FBP SUVR (log) | 0.371 | 0.357 | 0.002 | (0.139, 0.602) |
|  | YKL-40 (log) | -0.004 | -0.029 | 0.789 | (-0.032, 0.024) |
|  | GFAP (log) | 0.036 | 0.279 | 0.013 | (0.008, 0.064) |
| MTL Cortical Thickness (log) ~ | pTau-217 (log) | -0.112 | -0.345 | 0.001 | (-0.177,-0.047) |
| Hippocampal Volume (log) ~ | pTau-217 (log) | -0.278 | -0.254 | 0.013 | (-0.498, -0.058) |
| Retroactive Interference (log) ~ | MTL Cortical Thickness (log) | 0.471 | 0.121 | 0.217 | (-0.277, 1.218) |
|  | Hippocampal Volume (log) | 0.387 | 0.333 | 0.001 | (0.163, 0.611) |
|  | pTau-217 (log) | 0.182 | 0.143 | 0.202 | (-0.097, 0.46) |
| **Covariances** |  | *B* | Standardized Estimate | p-value | 95% CI |
| YKL-40 (log) ~~ GFAP (log) |  | 0.073 | 0.183 | 0.137 | (-0.023, 0.169) |

**Figure 2.**
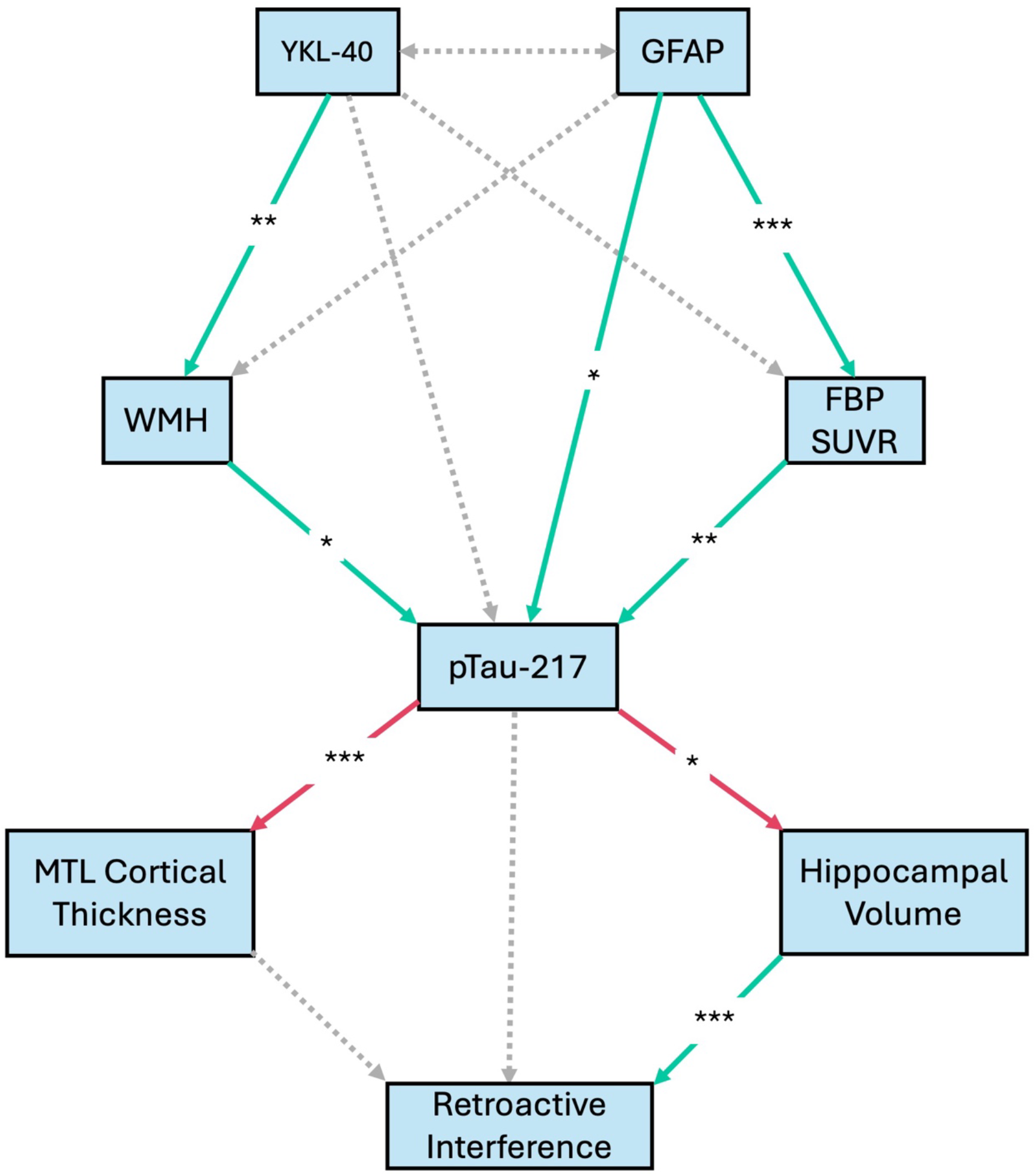
Path diagram Path diagram indicating pathway from neuroinflammatory markers (YKL-40 and GFAP) to downstream biomarkers and subsequent memory deficits. Solid lines demonstrate significant paths, and dotted lines represent non-significant paths. Green arrows represent positive associations, and red arrows represent negative associations. Paths indicated as significant (*) for p-values ≤ 0.05, (**) for p ≤ 0.01, and (***) for p ≤ 0.001. Results of path analyses are shown in Table 3.

**Figure 3.**
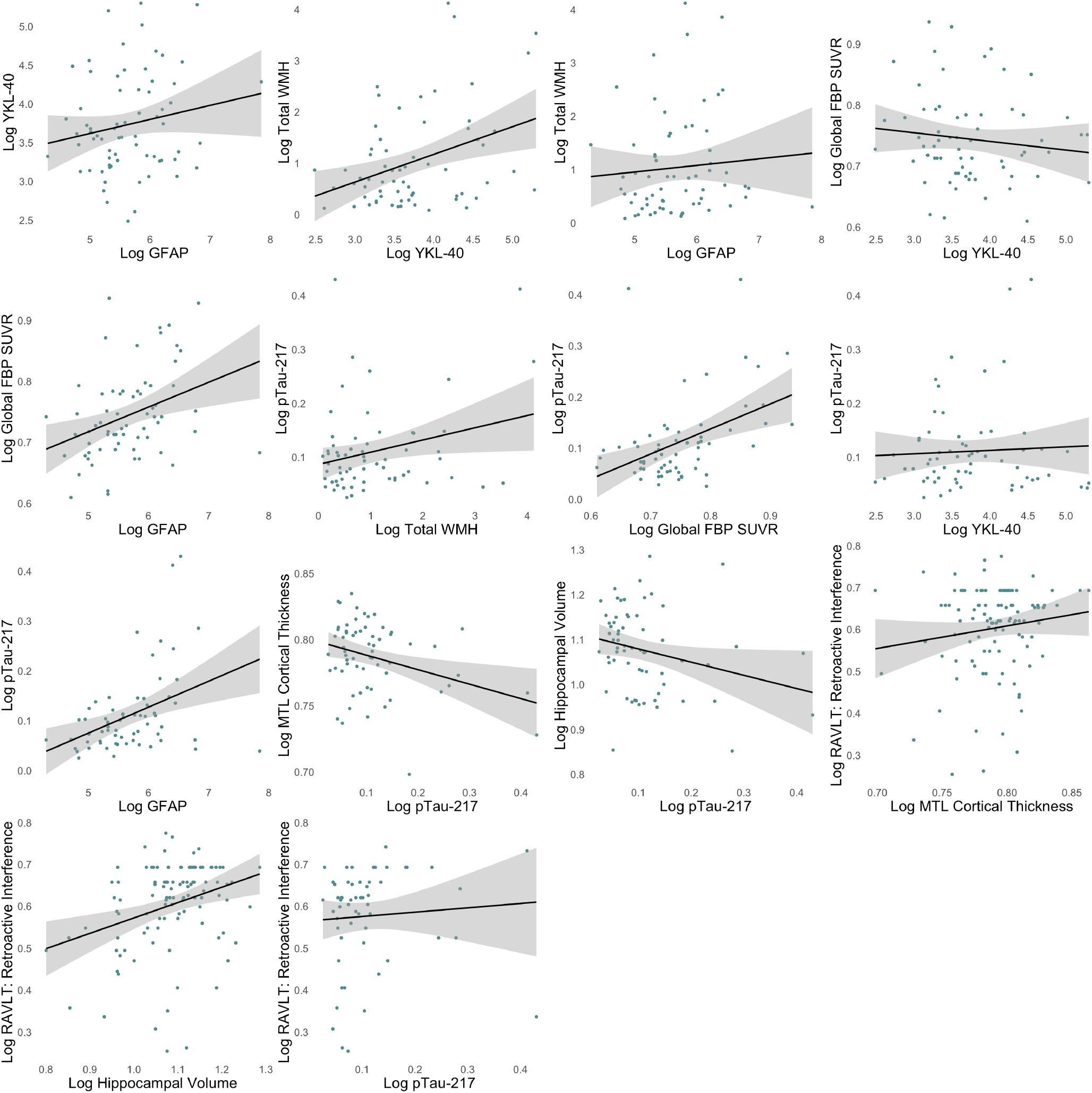
Simple Scatterplots. Scatterplots of associations among markers that were included in linear regressions, as part of the path analysis.

## 4. DISCUSSION

Our results support a model in which two glial cell activation and neuroinflammatory markers, YKL-40 and GFAP, are differentially related to small vessel cerebrovascular disease and amyloid, respectively, suggesting at least two pathways converging to phosphorylated tau. Phosphorylated tau was subsequently associated with neurodegeneration and lower memory performance in older adults. This pathophysiological cascade is depicted in Supplemental Figure 1.

While many studies showed a temporal ordering among amyloid, tau, and neurodegeneration ^45-48^, a similar temporal pathway has not been described with respect to cerebrovascular disease, tau, and neurodegeneration. However, previous work demonstrated associations between cerebrovascular disease and tau in both humans and mice ^32,33,49^. Similarly, prior work showed WMH are associated with MTL atrophy in both longitudinal and cross-sectional analyses ^30,31,50^. We did not explicitly model a relationship between amyloid and WMH, given that we did not expect a relationship between amyloid and WMH. Many previous studies reported no association between the two markers among clinically unimpaired individuals ^51,52^. Of interest, we expected to observe a direct effect of phosphorylated tau on memory given past studies suggesting alternative pathways toward memory loss ^53,54^, but we did not find a direct association between p-Tau217 and memory. It is possible that hippocampal volume fully mediates the association between phosphorylated tau and memory deficits.

We found that plasma YKL-40, a glycoprotein secreted and synthesized by astrocytes, microglia, and peripheral macrophages ^28,55^, was associated with WMH but not with Aβ. Previous work on serum YKL-40 demonstrated its association with WMH volume, and, further, that macroscopic and microscopic white matter damage mediated the correlation between serum YKL-40 and cognitive impairment in patients with small vessel disease ^28^.

In contrast to our findings with YKL-40, we found that plasma GFAP was associated with Aβ, but not with WMH. Prior work demonstrated that GFAP is associated with Aβ burden in unimpaired older adults and in those who are at risk for AD ^11,25,56^. Studies have also reported a link between GFAP and WMH ^27,56^; Shir et al. ^56^ demonstrated that the relationship between GFAP and WMH is only found in individuals with elevated levels of Aβ. Consistent with this understanding, the link between GFAP and WMH shown by Edwards et al. ^27^ was found in adults with Down syndrome, who also demonstrate elevated levels of amyloid ^57,58^.

Two previous studies did not show a link between GFAP and tau when Aβ is accounted for ^25,56^, but we found an association between GFAP and tau even when amyloid was included in the model. However, an important difference between their studies and ours is their use of tau-PET, which is more likely than plasma pTau-217 to reflect tau aggregation ^59^.

There are some limitations with the current study. One limitation is that path analysis can not examine bidirectional relationships that might be recursive in nature, for example between WMH and YKL-40, or between Aβ and GFAP. Due to the cross-sectional nature of the study design, we are unable to address the temporality or causality of the correlational associations found through the path analysis. With the limited parametric range of cognitive performance among clinically unimpaired older adults within our sample, we did not adjust for age in our path analysis model, as these associations are substantially driven by aging. Future work should include a more diverse sample of participants, as our study had predominantly non-Hispanic White participants. Another future direction is to include a broader array of markers of inflammatory disease and small vessel disease while applying a latent factor SEM approach. Given that plasma pTau-217 is associated with both amyloid and tau pathology ^59^, another future step would include using a more specific measure of tau aggregation, such as tau-PET. Regional distributions of amyloid and tau can also be considered with the use of PET, allowing for understanding more specific mechanistic roles of these two markers in this pathway.

In summary, we found that cerebrovascular injury and Aβ form two parallel and distinct pathways by which neuroinflammatory changes may alter Alzheimer’s pathological features, converging on the neurodegenerative cascade leading to cognitive decline. Identifying these pathways enables the early detection of biomarkers and the targeting of therapeutic interventions to the appropriate windows of disease. This understanding can also provide traction for addressing individual differences in AD vulnerability and resilience, potentially paving the way for personalized therapeutic strategies.

## Acknowledgments

We would like to thank the participants of the BEACoN study.

## Data Sharing Statement

Data will be shared upon request.

## Supplemental Results

**Supplemental Table 1.**
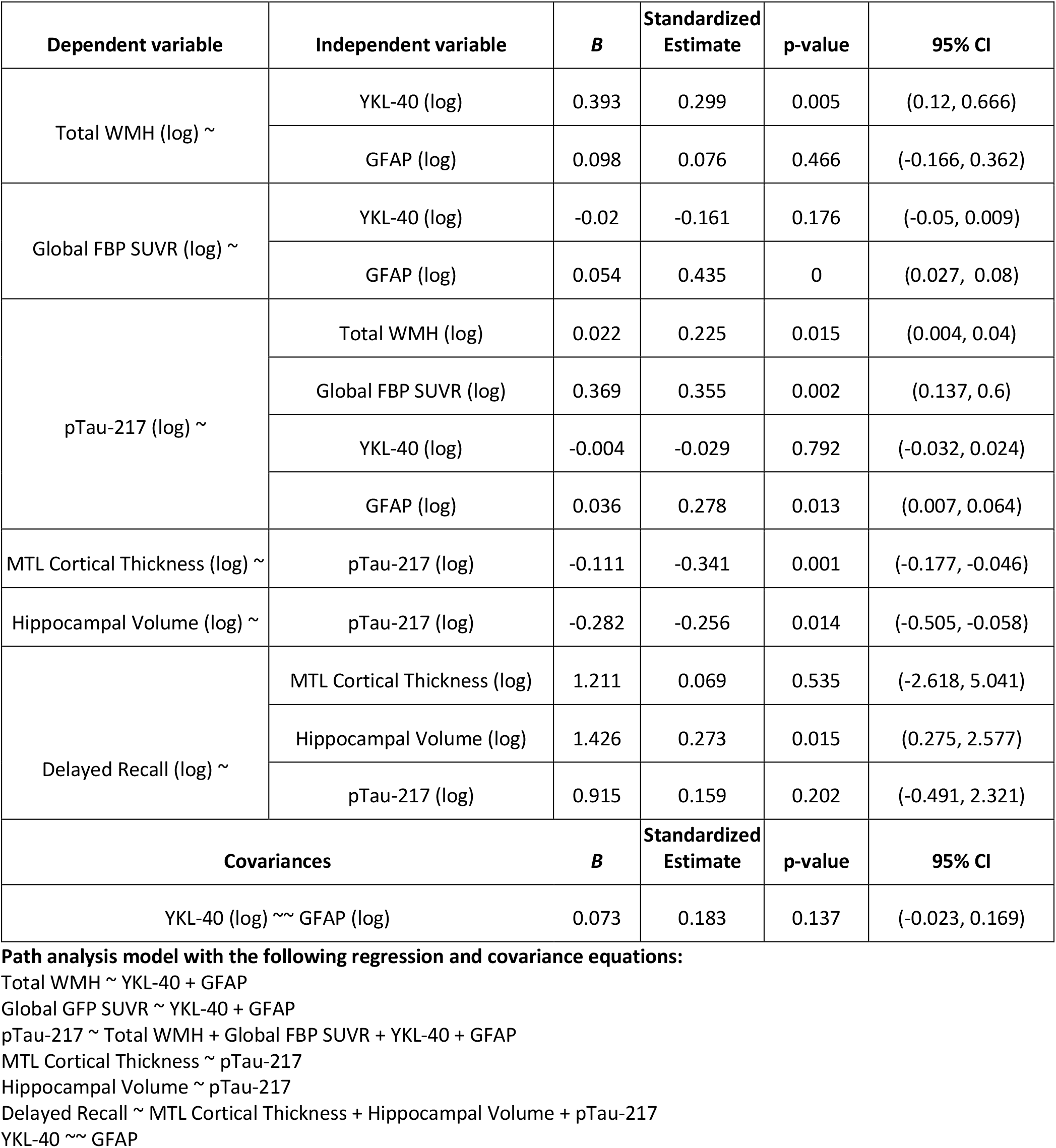

**Supplemental Figure 1.**
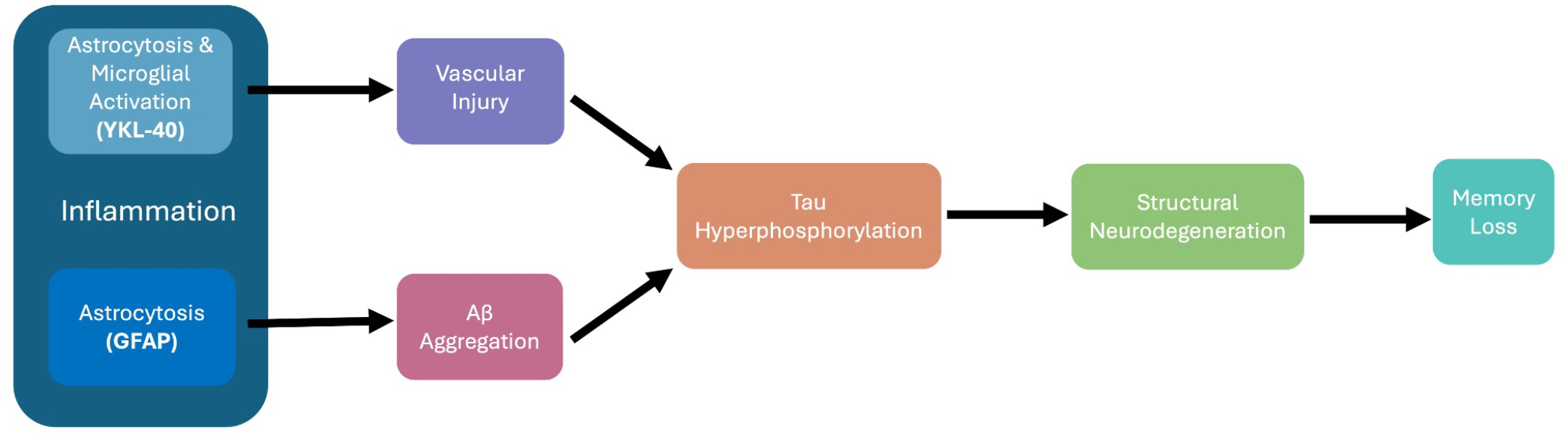
Conceptual mechanistic diagram. Conceptual diagram of the mechanistic cascade leading to memory deficits in older adults. Two parallel neuroinflammatory (plasma YKL-40 and GFAP) pathways highlight distinct mechanisms by which cerebrovascular injury (WMH) and Aβ (FBP SUVR) converge to tau pathology (plasma pTau-217), and subsequently to neurodegeneration (hippocampal and MTL cortical atrophy) and memory decline (RAVLT RI performance).

